# High fidelity sensory-evoked responses in neocortex after intravenous injection of genetically encoded calcium sensors

**DOI:** 10.1101/2023.03.09.531938

**Authors:** Austin Leikvoll, Prakash Kara

## Abstract

Two-photon imaging of genetically-encoded calcium indicators (GECIs) has traditionally relied on intracranial injections of adeno-associated virus (AAV) or transgenic animals to achieve expression. Intracranial injections require an invasive surgery and result in a relatively small volume of tissue labeling. Transgenic animals, although they can have brain-wide GECI expression, often express GECIs in only a small subset of neurons, may have abnormal behavioral phenotypes, and are currently limited to older generations of GECIs. Inspired by recent developments in the synthesis of AAVs that readily cross the blood brain barrier, we tested whether an alternative strategy of intravenously injecting AAV-PhP.eB is suitable for two-photon calcium imaging of neurons over many months after injection. We injected young (postnatal day 23 to 31) C57BL/6J mice with AAV-PhP.eB-Synapsin-jGCaMP7s via the retro-orbital sinus. After allowing 5 to 34 weeks for expression, we performed conventional and widefield two-photon imaging of layers 2/3, 4 and 5 of the primary visual cortex. We found reproducible trial-by-trial neural responses and tuning properties consistent with known feature selectivity in the visual cortex. Thus, intravenous injection of AAV-PhP.eB does not interfere with the normal processing in neural circuits. *In vivo* and histological images show no nuclear expression of jGCaMP7s for at least 34 weeks post-injection.

## 1 Introduction

Two-photon calcium imaging is a broadly used technique for recording neuronal population activity with cellular resolution *in vivo* (Grewe and Helmchen, 2009; Grienberger and Konnerth, 2012; Grienberger et al., 2022). Conventional two-photon microscopes record from relatively small fields of view (< 1 × 1 mm). The very first *in vivo* two-photon calcium imaging study from the neocortex imaged 100–200 neurons in small patches of neocortex (< 300 × 300 μm regions) (Stosiek et al., 2003). Imaging such small regions of neural tissue has yielded many insights into local microcircuit processing in various sensory systems including visual, olfactory, auditory, taste and somatosensory cortices (Ohki et al., 2005; Li et al., 2008; Kara and Boyd, 2009; Stettler and Axel, 2009; Bandyopadhyay et al., 2010; Chen et al., 2011a; Clancy et al., 2015). Incremental improvements in laser power, objective lens technology, scanning speed, and detector sensitivity have all contributed to an explosive growth in two-photon calcium imaging (Litvina et al., 2019; Abdelfattah et al., 2022). Moreover, the BRAIN Initiative has been a major driving force in creating a new class of optical instruments that can measure computation across large populations of neurons (with cellular resolution) over multiple cortical regions (Jorgenson et al., 2015). Large field of view two-photon mesoscopes can now image regions up to 5 × 5 mm with sub-cellular resolution (Sofroniew et al., 2016; Demas et al., 2021; Yu et al., 2021). Exceeding these advancements in instrumentation, innovations regarding the mode of introducing calcium sensors into neurons and the sensitivity of these sensors have been rather remarkable over the past 20 years.

The first calcium imaging studies *in vivo* relied on synthetic chemical indicator dyes like Oregon Green BAPTA-1 AM (OGB-1 AM) (Stosiek et al., 2003; Ohki et al., 2005). Then genetically encoded calcium indicators (GECIs), like GCaMP, were progressively refined for high signal-to-noise changes in fluorescence, rapid response kinetics and long-term labeling of neurons (Nakai et al., 2001; Palmer and Tsien, 2006; Tian et al., 2009; Chen et al., 2013). Initially, GECIs were introduced to the brain with intracranial injections of adeno-associated virus (AAV) carrying a GECI transgene. Then, numerous transgenic mouse lines for jGCaMP6 (Thy1, Emx1, Ai94, Ai162) and GCaMP7 (G7NG817, TRE-G) were created (Dana et al., 2014; Sato et al., 2015; Monai et al., 2016; Daigle et al., 2018).

Intracranial AAV injections are not ideal for large field of view calcium imaging. Most notably, intracranial injections using micropipettes label a relatively small volume of tissue, typically a sphere ~1–2 mm in diameter, so labeling large volumes of the cortex with this method requires AAV injections be made at multiple sites. Intracranial injections also create a gradient effect in labeling–neurons closer to the center of the injection site express more GECI and are thus brighter than neurons at the periphery (Tian et al., 2009; Chen et al., 2013; Bedbrook et al., 2018). Because local administration of AAV delivers a high copy number of viral genomes to each transduced cell, this strategy can result in excessive levels of GECI expression, causing GECIs to accumulate in cell nuclei and potentially leading to neuronal dysfunction (Tian et al., 2009; Chen et al., 2013; Resendez et al., 2016; Yang et al., 2018; Dana et al., 2019). Intracranial injections also have practical disadvantages—they are relatively time-consuming and invasive procedures, and they require extensive practice to perform without damaging the underlying neural tissue.

GECI-expressing transgenic animals have the obvious advantage of brain-wide labeling (Dana et al., 2014; Dana et al., 2018). However, there are potential caveats to this approach, as some GCaMP lines have abnormal behavioral phenotypes likely arising from aberrant neocortical activity (Steinmetz et al., 2017). Two-photon laser powers used for *in vivo* imaging in transgenic lines are rarely provided, but when cited, they appear to be much higher than the average powers used in experiments where intracranial injections are performed. For example, the Thy1 line needed 145 mW for imaging layer 2/3 neurons (100-250 μm depth) (Dana et al., 2014). The Emx1 and Ai94 required up to 70 mW for imaging layer 2/3 (Huang et al., 2021). These potentially damaging higher laser powers are likely due to dimmer neurons which reflect lower expression levels, compared to studies that used intracranial AAV injections. Finally, producing and validating new transgenic animal lines is a relatively slow process, limiting the speed at which new versions of GECIs can be incorporated into transgenic animals.

Virally-mediated, brain-wide expression of GECIs is now possible due to the development of AAV capsid variants capable of crossing the blood-brain barrier following intravenous injection (Deverman et al., 2016; Chan et al., 2017; Goertsen et al., 2022). The AAV-PhP.B capsid (Deverman et al., 2016) has already been used to achieve brain-wide GECI expression for functional imaging, but these studies required a high dose of virus (> 1×10^12^ genome copies per mouse) to achieve adequate expression. The newer AAV-PhP.eB capsid (Chan et al., 2017), an enhanced variant of AAV-PhP.B, is particularly promising for achieving brain-wide GECI expression because it appears to transduce the majority of neurons across cortical layers even with a substantially lower viral titer than its predecessor (Brown et al., 2021).

None of the previous studies using AAV-PhP.B or AAV-PhP.eB determined the reliability of neural responses over repeated trials (Allen et al., 2017; Hillier et al., 2017). In fact, one of the studies included “unreliable” cells that responded to only one stimulus repetition (Hillier et al., 2017). While behavioral modulation can influence trial-by-trial reliability of sensory responses (Stringer et al., 2019), the animals in this previous AAV-PhP.B study were anesthetized (Hillier et al., 2017). In an earlier study using synthetic calcium dyes, sensory-evoked trial-by-trial responses were found to be less reliable with Fluo-4 than OGB-1 (Bandyopadhyay et al., 2010).

Here, we show that intravenous injections of AAV-PhP.eB-Synapsin-jGCaMP7s is suitable for two-photon calcium imaging of sensory-evoked neural responses that have large amplitudes and extremely high trial-by-trial reliability. Moreover, while previous work assessed the viability of expression up to 10 weeks post injection, we demonstrate that this method provides stable jGCaMP7s expression for at least 34 weeks, with maintenance of robust calcium responses and no visible nuclear expression.

## 2 Materials and methods

### 2.1 Animals and AAV injections

All animal procedures were approved by the Institutional Animal Care and Use Committee at the University of Minnesota. Male and female C57BL/6J mice (n=13, postnatal age 23-31 days, mass 11-22 g; Jackson # 000664) were anesthetized with isoflurane (4-5% induction, 1-2% maintenance) and intravenously injected with 60-100 μl of sterile saline containing 4×10^11^ viral genomes of AAV-PHP.eB via the retro-orbital sinus of the left eye. Each AAV-PHP.eB capsid carried a plasmid for jGCaMP7s driven by the synapsin promoter and WPRE regulatory element (AAV-PHP.eB-Synapsin-jGCaMP7s-WPRE; Addgene # 104487-PHPeB). Virus injections took less than 10 minutes from the beginning of anesthesia induction to full recovery. Mice were imaged with two-photon microscopy 5 to 34 weeks after AAV injection.

### 2.2 Surgical procedures

Cranial windows were installed over the left primary visual cortex prior to two-photon imaging. Mice were initially anesthetized with a bolus injection of fentanyl citrate (0.05 mg kg^-1^), midazolam (5 mg kg^-1^), and dexmedetomidine (0.25 mg kg^-1^). During imaging, continuous intraperitoneal infusion with a lower concentration mixture (fentanyl citrate: 0.002–0.03 mg kg^-1^ hr^-1^, midazolam: 0.2–3.0 mg kg^-1^ hr^-1^, and dexmedetomidine: 0.010–0.15 mg kg^-1^ hr^-1^) was administered using a catheter connected to a syringe pump (O’Herron et al., 2020a). Heart and respiration rates, rectal temperature, and SpO2 levels were monitored throughout surgery and imaging. Lidocaine (0.02 mL, 2%) was injected subcutaneously before making incisions. A titanium headplate was attached to the skull using C&B Metabond dental cement (Parkell # S380). Craniectomies (3–4 mm in diameter) were made over the left primary visual cortex (V1), centered ~2.5 mm lateral to lambda and ~1.5 mm anterior to the transverse sinus. After performing durotomies over V1, the cranial windows were sealed with agarose (1.5% in artificial cerebrospinal fluid) and a glass coverslip (5 mm diameter, 0.15 mm thickness; Warner # D263).

### 2.3 Imaging

Two-photon imaging of the visual cortex was performed with microscope from Bruker coupled to an InSight X3 laser set at 940 nm (Spectra-Physics), as we have used previously (Liu et al., 2020; Cho et al., 2022). Imaging was performed with either a 10X objective lens (NA 0.5, ThorLabs # TL10X–2P) or a 25X objective lens (NA 1.05, Olympus # XLPLN25XWMP2). On both objectives, the correction collar was set to 0.15 mm to compensate for the aberration from the coverslip. Images acquired with the 10X lens were 1.6 mm × 1.6 mm to 1.9 mm × 1.9 mm in size and collected at 1024 × 1024 pixels so we could better resolve individual neurons. When imaging with the 10X lens, the integration time per pixel was set to 0.8 μs, and the frame period was 1.68 s. Images collected with the 25X objective were 500 μm × 500 μm to 535 μm × 535 μm in size and collected at 512 × 512 pixels. When imaging with the 25X lens, the integration time per pixel was set to 3.2–3.6 μs, and the frame periods ranged from 1.10-1.35 s.

### 2.4 Visual stimulation

Drifting square-wave grating stimuli (100% contrast, 1.5 Hz temporal frequency, and 0.033 cycles/degree spatial frequency) were presented on a 17-inch LCD monitor placed 15 cm from the mouse’s right eye. Blackout cloth was wrapped around the objective lens to prevent light from the visual stimulation monitor from entering the microscope’s photomultiplier tubes (O’Herron et al., 2012). Two types of visual stimuli were used, one to assess receptive field location (retinotopic mapping) and the other to measure orientation selectivity. The retinotopic mapping stimulus was presented in a 3 × 3 grid on the visual stimulation monitor (Shen et al., 2012; O’Herron et al., 2020b). The retinotopic stimulus consisted of vertically-oriented gratings drifting in one direction of motion within a circle occupying approximately one-ninth of the stimulus monitor. Drifting gratings were displayed in a sequence, starting with position 1 in the upper left of the screen. For determining orientation selectivity, the visual stimuli consisted of drifting gratings presented with 8 directions of motion in 45° steps. During functional imaging runs, visual stimuli were displayed for 5 imaging frames interleaved with 10 frames of blank (equiluminant uniform gray across the whole screen). Thus, grating stimuli were displayed for approximately 6 s preceded by approximately 12 s of blank. Each stimulus condition was repeated 3 times.

### 2.5 Histology

After imaging, mice were euthanized by intraperitoneal injection of pentobarbital (>180 mg kg^-1^) and transcardially perfused with 12 ml of chilled phosphate-buffered saline (PBS) followed by 12 ml of chilled 4% paraformaldehyde (PFA) in PBS. Brains were removed, placed in 4% PFA solution overnight at 4 °C, then transferred to PBS and stored at 4 °C. Coronal brain sections (50 μm thick) were cut using a vibratome (PELCO easiSlicer), mounted onto slides, dried overnight, and coverslipped with mountant (Invitrogen # P36980). Histological sections were imaged with two-photon microscopy using the Bruker microscope described above.

### 2.6 Data analysis

All images were analyzed with code written in MATLAB (MathWorks). Neuronal cell bodies were masked manually as follows. First the entire time series of TIFF images was averaged and then contrast optimized using adaptive histogram equalization. Donut shaped neurons across the entire field of view were then readily visible and could be manually selected, equivalent to using the ImageJ magic wand tool (Walker, 2014). Fluorescence time courses of each neuron F(t) were calculated by averaging over the pixels within each cell body mask. Only raw data were used to calculate time course and subsequent response statistics–no denoising was performed on our datasets. The response ΔF/F were computed as (F1 - F0)/ F0, where F1 was the average fluorescence across the entire stimulus window and F0 was the average fluorescence during the last approximately 6 s of the blank interval. When using retinotopic visual stimuli, significantly responsive neurons were defined by ANOVA across baseline and the nine retinotopic visual stimuli across three trials (*p* < 0.01). When using orientation and direction specific visual stimuli, significantly responsive neurons were defined by ANOVA across baseline and eight directions over three trials (*p* < 0.01). Trial-by-trial variability was measured as coefficient of variation (standard deviation/mean) for visually responsive neurons. For each neuron, the coefficient of variation was calculated for the response to its preferred stimulus. Thus, during retinotopic mapping runs, the single stimulus location (of the nine presented) which gave the largest mean response over three trials was used to calculate trial-by-trial variability. Similarly, for orientation/direction mapping runs, the single stimulus direction (of the eight presented) which produced the largest mean response was used to determine trial-by-trial variability.

## 3 Results

### 3.1 Expression pattern across cortical layers following intravenous injection of AAV GCaMP7

Five to 34 weeks after intravenous injection, we examined jGCaMP7s expression patterns with *in vivo* two-photon microscopy or postmortem histological samples in thirteen mice (e.g., see Fig. 1). There are very few bona fide neurons in layer 1 and thus, as expected, little GCaMP7 expression was seen in layer 1. In contrast, robust expression was present in cortical layer 2/3. In previous work, we showed that intracranial injection of AAV.hSyn.GCaMP6s produces weak expression in cortical layer 4 (Liu et al., 2020) and thus we had resorted to OGB-1 AM dye for layer 4 calcium imaging (O’Herron et al., 2020a). Here we show that with AAV-PhP.eB-Synapsin-jGCaMP7s, there is no “gap” in expression in layer 4 when imaging *in vivo* (Fig. 1B, Supplementary Video 1) and when examining coronal histological sections (Fig. 1C–D). *In vivo,* we exceeded the 450 μm two-photon depth limit for functional imaging of fluorophores when they are approximately homogeneously labeled across cortical layers (Takasaki et al., 2020). Thus, our *in vivo* imaging included layer 5 pyramidal cell bodies at 550 μm (see Fig. 1). We could not image layer 6 neurons *in vivo* with two-photon microscopy, but layer 6 neurons are clearly labeled with this method as evident in histology (Fig. 1C–D). In our histological samples, we also saw jGCaMP7s expression in multiple brain regions below the neocortex (e.g., see Fig. 1C–D), but here we have focused on the expression pattern of AAV GCaMP7 specifically in V1, from several weeks post-injection of the AAV through more than eight months post-injection. We saw no nuclear expression even at 16-34 weeks after injection of AAV-PhP.eB (Fig. 1D and Supplementary Video 2).

**Figure 1.**
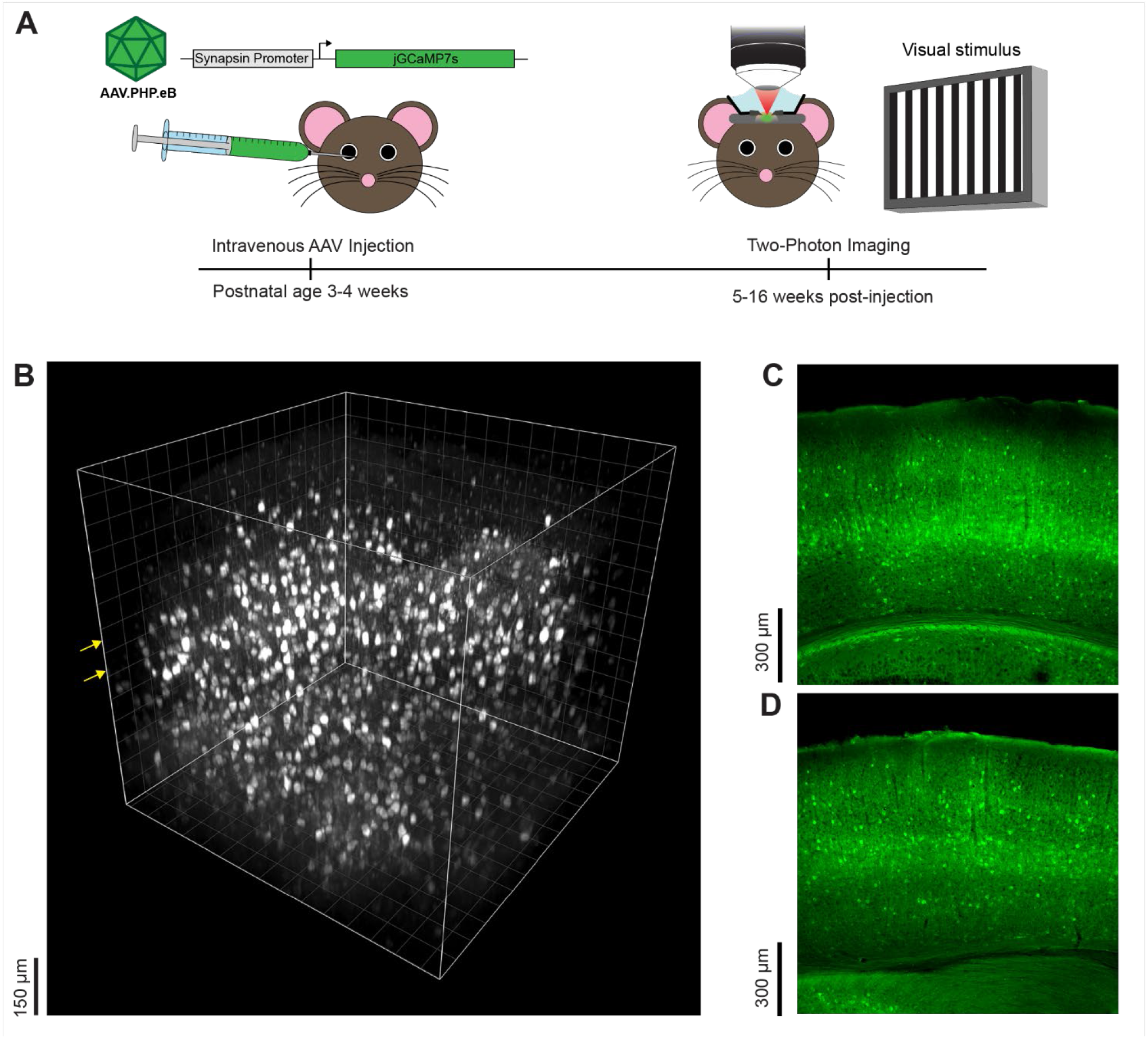
Experimental approach and representative cortical expression pattern following intravenous injection of AAV GCaMP7. **(A)** Timeline and overview of experimental methods. Three- to four-week-old mice were intravenously injected with AAV-PhP.eB-Synapsin-jGCaMP7s via the retro-orbital sinus. Five to thirty-four weeks later, sensory-evoked neural responses in the visual cortex were recorded with two-photon microscopy and the tissue was thereafter preserved for postmortem histology. **(B)** Side view of a three-dimensional volume of GCaMP7 expressing neurons obtained *in vivo,* 16 weeks after intravenous injection of the AAV-PhP.eB. The *x-y-z* dimensions of the volume are 750 μm × 750 μm × 700 μm and the first z plane is 50 μm above the pia. The grid line interval is also 50 μm. Yellow arrows indicate the planes of imaging where visually-evoked responses were recorded (see Figure 4). **(C)** Representative histological coronal section of the mouse visual cortex 8 weeks after intravenous injection of AAV-PhP.eB shows neural expression across cortical and sub-cortical layers. **(D)** Histological coronal section taken 34 weeks after intravenous injection of AAV-PhP.eB.

We next examined the fidelity of trial-by-trial responses in jGCaMP7s-expressing neurons *in vivo* using two different types of visual stimuli (retinotopy and orientation) in a subset of animals.

### 3.2 Reliable trial-by-trial neural responses to retinotopic visual stimuli after intravenous injection of AAV GCaMP7

Varying the location of the visual stimulus activated spatially distinct clusters of neurons in mouse V1 (*n* = 7 mice). Specific neurons responded optimally to a single location over repeated stimulus presentations (see Fig. 2). The shift in retinotopic location across the field of view (Fig. 2B) matched the known organization of the coarse retinotopic map present in mouse V1 (Schuett et al., 2002; Zhuang et al., 2017). Specifically, neurons located rostrally (anteriorly) responded to low elevation visual stimuli, e.g., in Fig. 2 neuron 1 responds to visual stimulus position 7 (bottom left of monitor). Neurons located caudally (posteriorly) responded to high elevation stimuli, e.g., in Fig. 2, neuron 8 responded to stimulus position 1 (top left monitor). Neurons located centrally in V1 (and thus in the middle of the field of view of imaging) responded to medium elevation visual stimuli, e.g., in Fig. 2, neurons 4 and 6 respond to stimulus position 4, which is directly between stimulus positions 1 and 7 (see Fig. 2B). Most importantly, the trial-by-trial variability of GCaMP7 neural responses to retinotopic visual stimuli was low. In the large region imaged shown in Fig. 2, the median coefficient of variation was 0.46 across all responsive neurons (see Fig. 2D). In mouse V1 studies, typically full-field visual stimuli that vary in direction and orientation are presented. Therefore, we next examined the trial-by-trial reliability of neural responses to stimuli varying in direction and orientation.

**Figure 2.**
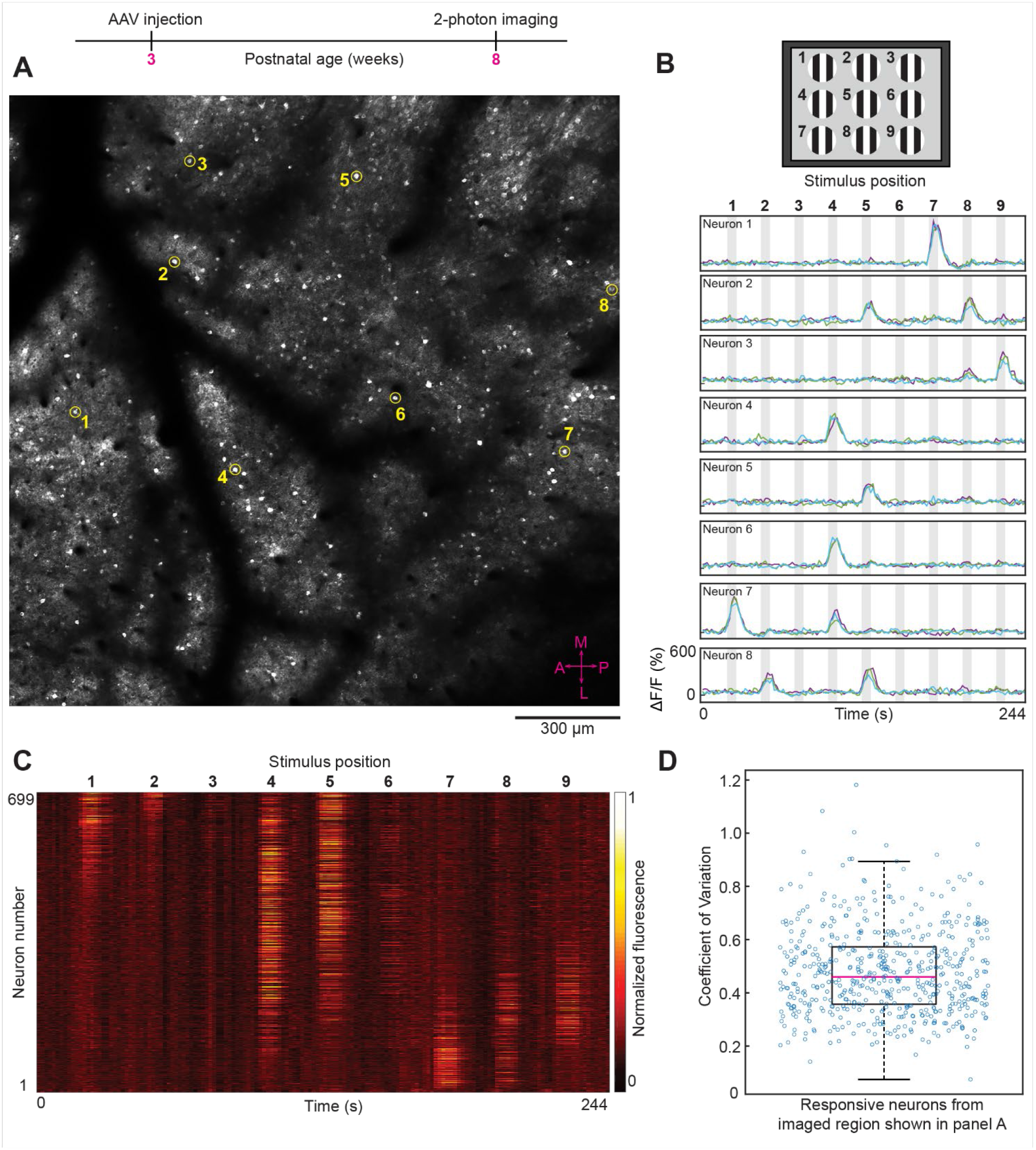
Two-photon imaging of neural responses to retinotopic (position dependent) visual stimuli after intravenous injection of AAV GCaMP7. **(A)** *In vivo* image from layer 2/3 (imaging plane 206 μm below pia) of a large region (1740 × 1740 μm) from mouse visual cortex. The average laser power used was 50 mW with a ThorLabs 10× objective lens. The eight circled neurons (numbered 1–8) were randomly selected to show their visually-evoked responses (see panel B). The compass in the bottom right corner (shown in magenta color) represents the medial, lateral, anterior and posterior coordinates of the imaged region. **(B)** Top panel, schematic of the visual stimulus sequence used for retinotopic mapping. Small patches of drifting gratings were sequentially presented in nine locations within a 3 × 3 grid, starting with position 1 and ending with position 9. Bottom panel, time courses for the eight example neurons (numbered 1–8) shown in panel A. Gray shaded areas in the time course plots correspond to periods of visual stimulation. The bolded numbers above each shaded area represent the stimulus position. Fluorescence traces from three trials are superimposed for each neuron. X and Y axes scales are identical for all eight examples shown. **(C)** Heat map of calcium responses from all 699 neurons that were masked in the imaged region shown in panel A. Each horizontal line represents a single neuron trace (response averaged across three trials). **(D)** Box plot showing the visually-evoked trial-by-trial variability (measured as coefficient of variation) for the 551 neurons that were significantly responsive to visual stimuli (*p* < 0.01 ANOVA across baseline and the 9 retinotopic visual stimuli across three trials). For the box plots in this and subsequent figures, the middle red line represents the median, the box represents the 25% and 75% quartiles, and the whiskers represent the 1 and 99 percentiles.

### 3.3 Reliable trial-by-trial neural responses to orientation selective visual stimuli after intravenous injection of AAV GCaMP7

In all field of views examined, varying the direction and orientation of visual stimuli produced one population of highly tuned orientation-selective neurons and another population of untuned GCaMP7 neurons (*n* = 10 mice). Tuned and untuned neurons were always interspersed in the same imaging field of view. For example, compare neurons 1 and 2 in Fig. 3A–C; neurons 3 and 4 in Fig. 3E–G and neurons 5 and 6 in Fig. 3I–K. Previous studies using other types of calcium sensors, e.g., OGB-1 AM, established two populations of neurons, tuned excitatory neurons and untuned inhibitory neurons (Sohya et al., 2007; Kerlin et al., 2010). Since we wanted to examine the fidelity of intravenous injection of AAV GCaMP7, without potential interference from other reagents, we did not attempt to tag the various classes of inhibitory neurons. AAV-PhP.eB has already been shown to transduce excitatory neurons and the three major classes of inhibitory neurons (Brown et al., 2021). Most critically, orientation and direction specific stimuli evoked GCaMP7 neural responses with very low trial-by-trial variability. In the imaged regions shown in Fig. 3, the median coefficient of variation was 0.59 in layer 2/3 (Fig. 3D), 0.55 in layer 4 (Fig. 3H) and 0.48 in layer 5 neurons (Fig. 3L). Intracranial injections of AAV typically result in viable expression and functional imaging for 1–2 months post injection. Thereafter nuclear expression and attenuation of sensory-evoked responses have been observed (Tian et al., 2009; Chen et al., 2013). Therefore, we next determined if intravenous AAV injections can maintain normal expression and high trial-by-trial reliability of responses for several months post injection (see below and Fig. 4, Supplementary Fig. 1).

**Figure 3.**
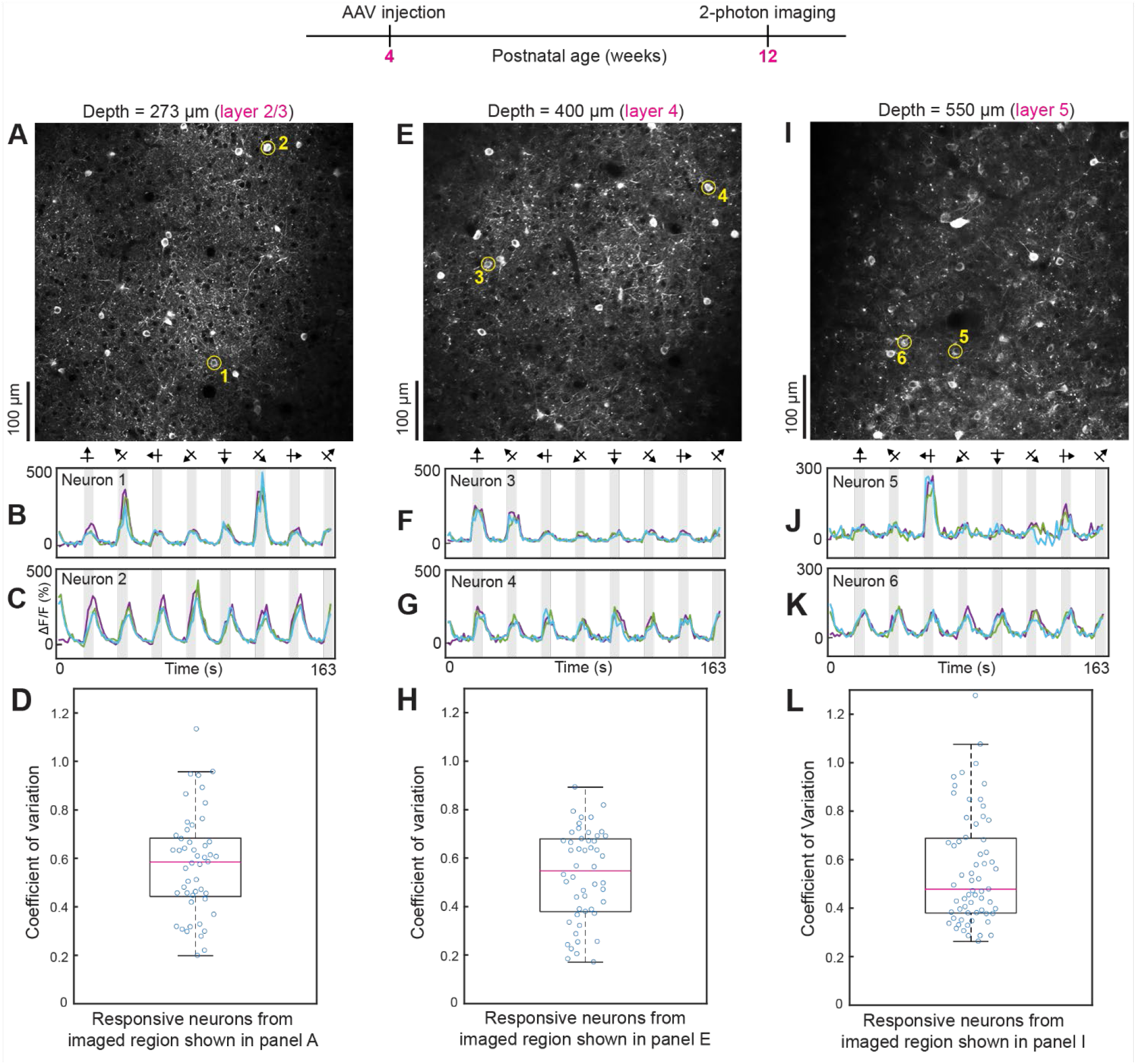
Two-photon imaging of neural responses to orientation and direction selective visual stimuli after intravenous injection of AAV GCaMP7. (**A**–**D**) Data from layer 2/3 (imaging plane 273 μm below the pia). The average laser power used was 30 mW. For the data shown in this and subsequent figures, a high NA 25x Olympus objective lens was used. The two circled neurons in panel A (numbered 1–2) were selected to show tuned and untuned visual responses, respectively (see time courses in panels B, C). For all time courses shown, three trials are overlaid. Panel D shows a box plot of the coefficient of variation of calcium responses for neurons that were significantly responsive to visual stimuli (*p* < 0.01, ANOVA across baseline and eight directions over three trials). (**E**–**H**) Data from layer 4 (imaging plane 400 μm below the pia). The average laser power used was 48 mW. Conventions for time courses and box plots are the same as shown in previous panels. (**I**–**L**) Data from layer 5 (imaging plane 550 μm below the pia). Average laser power during imaging was 56 mW.

**Figure 4.**
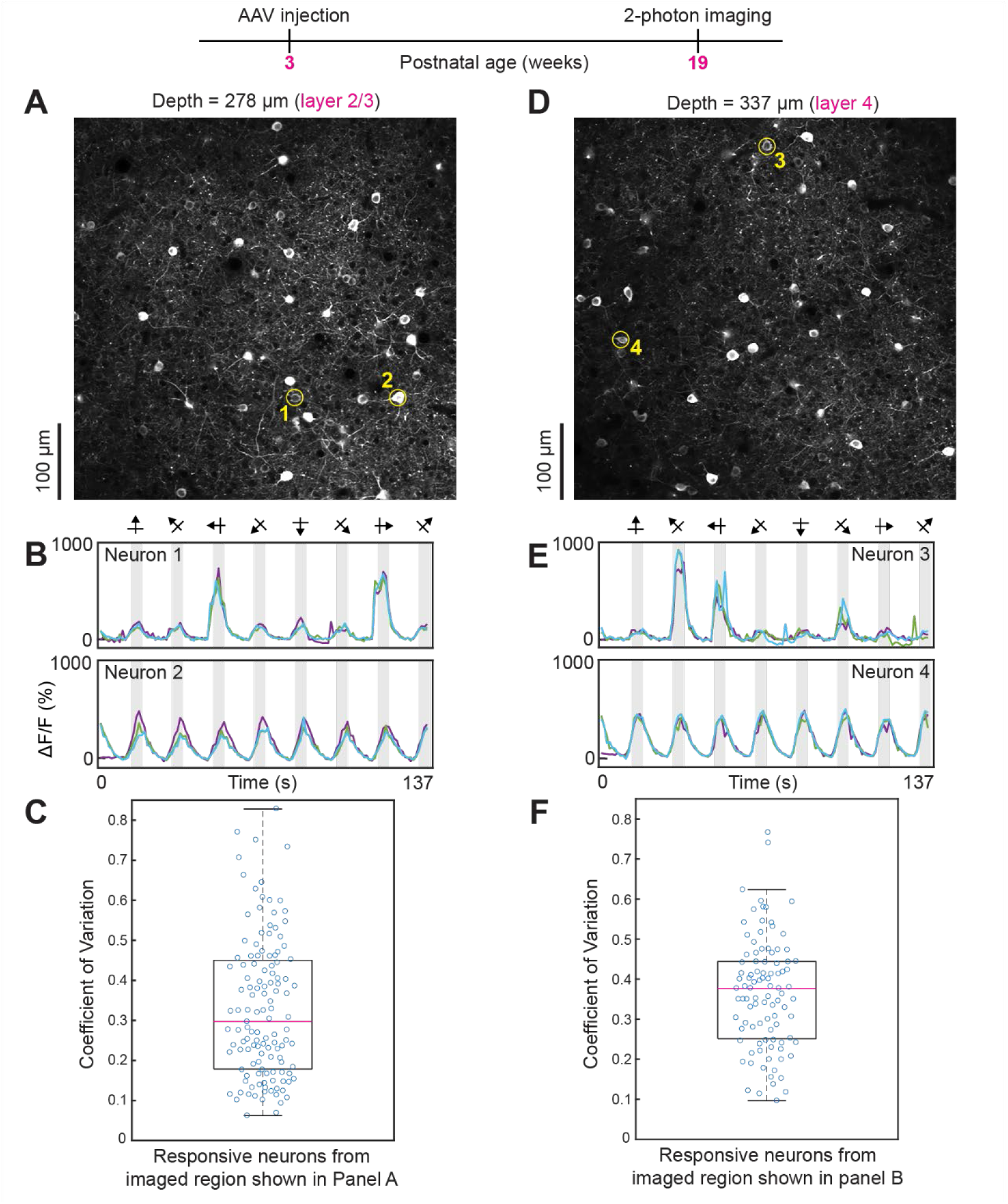
Robust visual responses persist even after 16 weeks of intravenous injection of AAV GCaMP7. **(A–C)** Data from layer 2/3. The average laser power during imaging was 26 mW. **(D-F)** Data from layer 4. The average power used was 32 mW. Conventions for two-photon imaging planes, time courses and box plots, as in previous figures.

### 3.4 Robust visual responses persist for many months after intravenous injection of GCaMP7

The distribution of animals imaged *in vivo* across post-injection periods was as follows. Five weeks after intravenous injection *n* = 1 mouse, six weeks after injection *n* = 3 mice, 7 weeks post injection *n* = 1 mouse, 8 weeks post injection *n* = 2 mice, 9 weeks post injection *n* = 1 mouse, 10 weeks post injection *n* = 1 mouse, 11 weeks post injection *n* = 1 mouse, 16 weeks post injection *n* = 1 mouse and 34 weeks post injection *n* = 1 mouse. Response amplitudes were typically >300% across all periods and all cortical layers examined (see Figs. 2–4, Supplementary Fig. 1 and Supplementary Video 2). The trial-by-trial variability of responses to visual stimuli (measured as the coefficient of variation) was consistently below 0.6 across all imaging periods (5–34 weeks post injection) and in all cortical layers examined (layers 2/3, layer 4 and layer 5)(see Figs. 2–4 and Supplementary Fig. 1).

## 4 Discussion

### 4.1 Reliable trial-by-trial feature selective responses with intravenous injection of AAV GCaMP7

We have shown that intravenous injection of AAV-PhP.eB-Synapsin-jGCaMP7s provides robust and stable expression for high fidelity two-photon imaging of sensory-evoked neural responses in mouse visual cortex. The widespread expression is sufficiently strong for conventional small field of view two-photon imaging and wide-field two-photon imaging. Feature selectivity such as retinotopy and orientation-selectivity was preserved with intravenous injection of AAV GCaMP7.

Most importantly, trial-by-trial sensory-evoked responses were highly reproducible in these intravenous AAV GCaMP7 experiments in mice, reminiscent of prior work using single-unit recordings from the cat visual cortex (Kara et al., 2000). While animals engaged in behavioral tasks may evoke eye movements and attention modulation that correlate with trial-by-trial variability (Stringer et al., 2019), our animals were anesthetized. Prior work in macaque monkeys has shown that when subjects maintain fixation, trial-by-trial variability of visually-evoked responses is markedly lower during the awake state compared to during anesthesia (Festa et al., 2021). Thus, it is not a foregone conclusion that anesthesia necessitates reliable sensory-evoked responses over repeated trials. For example, in the auditory and visual cortex, there have been reports of highly variable trial-by-trial responses to sensory stimulation during two-photon calcium imaging in anesthetized mice (Bandyopadhyay et al., 2010; Jia et al., 2010; Chen et al., 2011b). The type of anesthesia, the experimental preparation quality and sensor performance can all heavily impact the fidelity of responses (Bandyopadhyay et al., 2010; Grienberger et al., 2022). Even in visual cortical brain slices, monosynaptic activation of spiny stellate neurons in layer 4 can evoke responses with a wide range of trial-by-trial variability (Stratford et al., 1996). For example, the coefficient of variation of responses in layer 4 neurons is nearly an order of magnitude lower when stimulating LGN afferents compared to when stimulating layer 5/6 neurons (Stratford et al., 1996).

### 4.2 Prior use of intravenous injection of GCaMP6, GCaMP7 and GCaMP8

Previous studies have used AAV-PhP.eB-Syn-GCaMP6s for widefield one-photon imaging in neocortex (Michelson et al., 2019; Xiao et al., 2021). Additional work used AAV-PhP.eB-CAG-FLEX-jGCaMP7s in Cre-driver mice to achieve GECI expression in specific cortical layers and then used 2-photon imaging to measure the effects of anesthesia on spontaneous neuronal activity (Bharioke et al., 2022). However, to our knowledge no previous studies have demonstrated retinotopic and orientation-selective responses at the single-neuron level or the high-fidelity trial-to-trial responses achievable with intravenous injections of AAV-PhP.eB-Synapsin-GCaMP7s. Even the most recent soma-targeting AAV-PhP.eB-GCaMP8 study examined visual cortical responses to orientation and direction selective stimuli by intracranial injection of the AAV, not intravenous injection (Grodem et al., 2023).

### 4.3 Closing the layer 4 gap with intravenous injection of GCaMP7

Unlike previous work showing that intracranial injection of AAV2/9.hSyn.GCaMP6s.WPRE.SV40 produces very weak expression in cortical layer 4 (Liu et al., 2020), here we show that with intravenous injection of AAV-PhP.eB-Synapsin-jGCaMP7s, there is no “gap” in layer 4 expression.

### 4.4 Viable expression and high-fidelity sensory responses persist for many months after intravenous injection of GCaMP7

In all past studies examining AAV-PhP.B- and AAV-PhP.eB-mediated GCaMP, the maximum post-injection period examined was 10 weeks (Allen et al., 2017; Hillier et al., 2017; Michelson et al., 2019; Brown et al., 2021; Grodem et al., 2023). We have extended this timeframe by demonstrating that expression remains stable and free of nuclear labeling for at least 34 weeks after transduction. Moreover, at this age, trial-by-trial responses had low variability, just as we found at earlier time points.

### 4.5 Future directions

Intravenous injection of AAV GCaMP7 appears to be suitable for long-term chronic two-photon imaging across multiple brain regions in projects that involve learning and memory, recovery from experience-dependent plasticity, aging and Alzheimer’s disease. The 34 week post-injection example we show (Supplementary Fig. 1) is from a 37-week-old mouse, which is considered “middle-age” (Flurkey et al., 2007). It is possible that quality expression and reliable responses will be present even in “old age” mice (>72 weeks).

For any of the long-term imaging paradigms mentioned above, a key advantage of AAV-mediated GECI expression over transgenic animals is flexibility. To change the transgene carried by an AAV capsid is straightforward, whereas creating new transgenic animal lines is a difficult and lengthy process. This flexibility is especially advantageous for calcium imaging because new generations of GECIs with enhanced performance and cell-type specific promoters are regularly engineered (Grodem et al., 2023). Additionally, intravenous AAV injections could be used to label neurons in other transgenic lines, e.g., disease models (as noted above) without the need for crossbreeding with a transgenic GECI line.

Finally, it is important to note that the AAV-PhP.eB capsid can efficiently cross the blood brain barrier only in a subset of mouse strains, such as C57BL/6J, which have particular isoforms of the Ly6a receptor (Hordeaux et al., 2019; Huang et al., 2019; Batista et al., 2020). However, several new AAV variants cross the blood brain barrier in all mouse strains and even in non-rodent species, including non-human primates (Ravindra Kumar et al., 2020; Goertsen et al., 2022). It is expected that refinement of these and other newly developed tools, such as soma- and axon-targeted expression of GCaMP (Grodem et al., 2023), will make two-photon calcium imaging in all mammalian species completely ‘plug-and-play’ in the near future.

## 6 Author Contributions

AL and PK designed experiments. AL collected data. AL and PK performed data analysis and wrote the manuscript. Both authors reviewed and approved the final submitted manuscript.

## 8 Acknowledgments

We thank D. Farinella and H. Jayakumar for optics support in the Kara laboratory, Viviana Gradinaru at CalTech for developing the PhP.eB technology and donating it to Addgene. We also thank Aaron Kerlin, Madhu Kannan and Zane Crabtree for comments on the manuscript.

## 9 Data Availability Statement

The raw data supporting the conclusions of this article will be made available by the authors, without undue reservation.

---

**Supplementary Figure 1.**
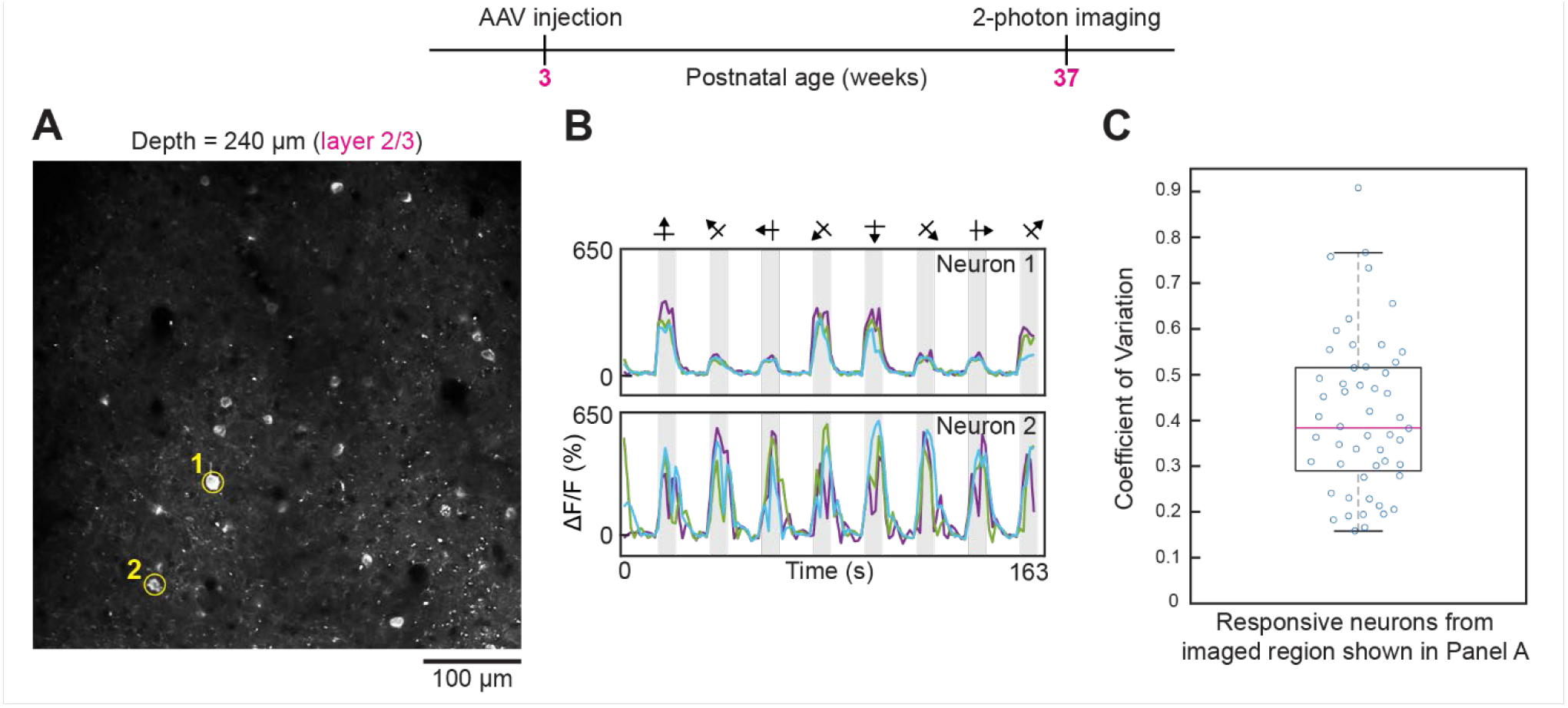
Two-photon imaging of neural responses 34 weeks after intravenous injection of AAV GCaMP7. (**A-B**) Data from layer 2/3 (imaging plane 240 μm below the pia). The average laser power used was 45 mW. The two circled neurons in panel A (numbered 1–2) were selected to show tuned and untuned visual responses, respectively (see time courses in panel B). For all time courses shown, three trials are overlaid. (**C**). Box plot of the coefficient of variation of calcium responses for neurons that were significantly responsive to visual stimuli (*p* < 0.01, ANOVA across baseline and eight directions over three trials).

## See supplementary material for links to supplementary videos

**Supplementary video 1.** *In vivo* z-stack demonstrating cortical expression following intravenous injection of AAV GCaMP7. Rotating side perspective of the 3D volume reconstruction shown in Figure 1.

**Supplementary video 2**. Raw two-photon imaging frames showing GCaMP7 responses in layer 2/3 of mouse V1 *in vivo*. Raw data frames (left) and frame-locked presentation (right) of blank (gray) and drifting grating visual stimuli. Data are shown for 25 imaging frames: 10 frames blank, followed by 5 frames of visual stimulation and then 10 frames blank. The duration of each frame was 1.14 s. The entire x-y field of view shown in the 2 two-photon imaging frames represents a region of 500 μm × 500 μm. Movie corresponds to data shown in Fig. 4D–F (16 weeks after intravenous injection).

